# GeneTerpret: a customizable multilayer approach to genomic variant prioritization and interpretation

**DOI:** 10.1101/2020.12.04.408336

**Authors:** Roozbeh Manshaei, Sean DeLong, Veronica Andric, Esha Joshi, John B. A. Okello, Priya Dhir, Kirsten M. Farncombe, Kelsey Kalbfleisch, Cherith Somerville, Rebekah K. Jobling, Stephen W. Scherer, Raymond H. Kim, S. Mohsen Hosseini

## Abstract

Variant interpretation is the main bottleneck in medical genomic sequencing efforts. This usually involves genome analysts manually scouring through a multitude of independent databases, often with the aid of several and mostly independent computational tools.

To streamline the variant interpretation process, we developed *GeneTerpret* platform that collates data from current interpretation tools and databases, and applies a phenotype-driven query to categorize the variants identified in a given genome. The platform assigns quantitative validity scores to genes by query and assembly of the current genotype-phenotype data, sequence homology, molecular interactions, expression data, and animal models. The platform uses the American College of Medical Genetics (ACMG) criteria to categorize variants into five tiers (from benign to pathogenic). The platform then outputs a prioritized list of potentially causal variants/genes in a given genome for a specific case.

*GeneTerpret* is a flexible and free platform designed to streamline the variant interpretation process through a unique interface, with improved ease, speed and accuracy. This unique integrated system provides effective validity and pathogenicity modules to assess genetic variant data and allows the user to decide which output and impact level should be considered in this process. The platform can be accessed and used online at https://geneterpret.com.

## INTRODUCTION

Rapid advances in DNA sequencing technologies have enabled the revolutionary use of clinical genomic data to support precision medicine initiatives, improving patient care and medical management. The aim of Genomic Variant Interpretation (GVI) is to identify one or a few medically relevant variants from hundreds of thousands in a given genome.^1^ To do this accurately, the genomic evidence supporting the association of a candidate gene with a disease of interest (gene-disease validity) and the detrimental effect of a variant on the gene function (variant pathogenicity) must be well evaluated. Although a multitude of independent computer programs is available to aid the GVI process, GVI routinely requires manual interpretation by a human analyst who leverages expertise, insight and phenotypic knowledge to curate a list of candidate variants. As such, the process is often tedious, repetitive, time-consuming, and may be prone to human errors. It is therefore not surprising that discordance exists among germ-line variant classifications across labs, diseases and variant types.^2^ Part of the discordance is due to different technologies and variant interpretation pipelines. So, a unifying platform for GVI is much needed.

The currently available GVI platforms and tools, although helpful, usually lack proper validation, and often lack an interactive comprehensive interpretation tool. A growing number of these tools have been bundled into commercial or free packages to aid in genome interpretations. Some of the most commonly available commercial packages are VarSome^3^, NAVIFY^4^, QIAGEN Clinical Insights (QCI)^5^, and CureMatch Bionov^6^. The most widely-used freely available packages include Exomiser^7^, Genetic Variant Interpretation Tool (GVIT)^8^, InterVar^9^, and CharGer^10^. A common feature among these packages is their ability to assess individual variant pathogenicity using the American College of Medical Genetics (ACMG) criteria. However, these tools usually ignore the gene-disease validity part of GVI, and in their application of ACMG criteria, manual adjustments by the human interpreter are required. Moreover, these existing platforms lack the flexibility to provide an iterative reweighting workspace for the user to define what evidence/terms should be considered, and how to execute these evidence/terms sufficiently.

To address the need for a customizable GVI platform, we developed *GeneTerpret* as a visual analytics tool to accelerate prioritization with an easy interface for expert interaction. The platform considers both phenotypic and genomic information to produce and prioritize a list of putative medically relevant variants. The platform can accurately analyze genomes from singletons, trios or entire cohorts, and can extract a significantly more manageable candidate-gene list for a human analyst to review. Overall, *GeneTerpret* improves the GVI process by increasing the speed and potentially reducing associated costs, while providing the analyst with the freedom to customize the platform’s parameters, filters and outputs. By adopting the use of a platform as *GeneTerpret* across labs, we postulate that its use could also help reduce the number of inter-lab variability in variant interpretation accuracy.

## MATERIALS AND METHODS

### GeneTerpret Workflow and Implementation

The *GeneTerpret* platform execution is modular and customizable, allowing the user to generate candidate gene lists based on different inputs and parameters, such as specific tissue type, phenotype, and known gene(s). It accepts genotype data and family information in Variant Call Format (VCF) and Pedigree (PED) file format. The outputs of the *GeneTerpret* analysis can be a more refined list of genes, associated phenotype(s) and VCF files for further consideration. An overview of the *GeneTerpret* platform workflow is summarized in Figure 1. More details on the backend and web implementation are presented in the supplementary file (S1 section). In brief, the workspace area is accessible through a graphic user interface (GUI) and acts as a prototypical canvas upon which phases of query and data processing are performed. Each entry is represented as a node that can be flexibly added or removed to achieve a user-desired analysis scheme. Nodes can be intuitively connected for data or modules of compatible inputs and output to allow the flow of data, with the platform and associated algorithms executing the corresponding module functions in the backend. This way, the user can quickly apply these complex functions to data, triggering the execution of the backend functionalities. Once the results are ready, the user can download the compressed file of prioritized genes for review, while the inputted data and settings from a recent analysis session remain open for the user to re-customize and fine-tune the analysis. Supplementary Figure S1 shows an example scenario of the GUI during the implementation of the platform workflow.

**Figure 1).**
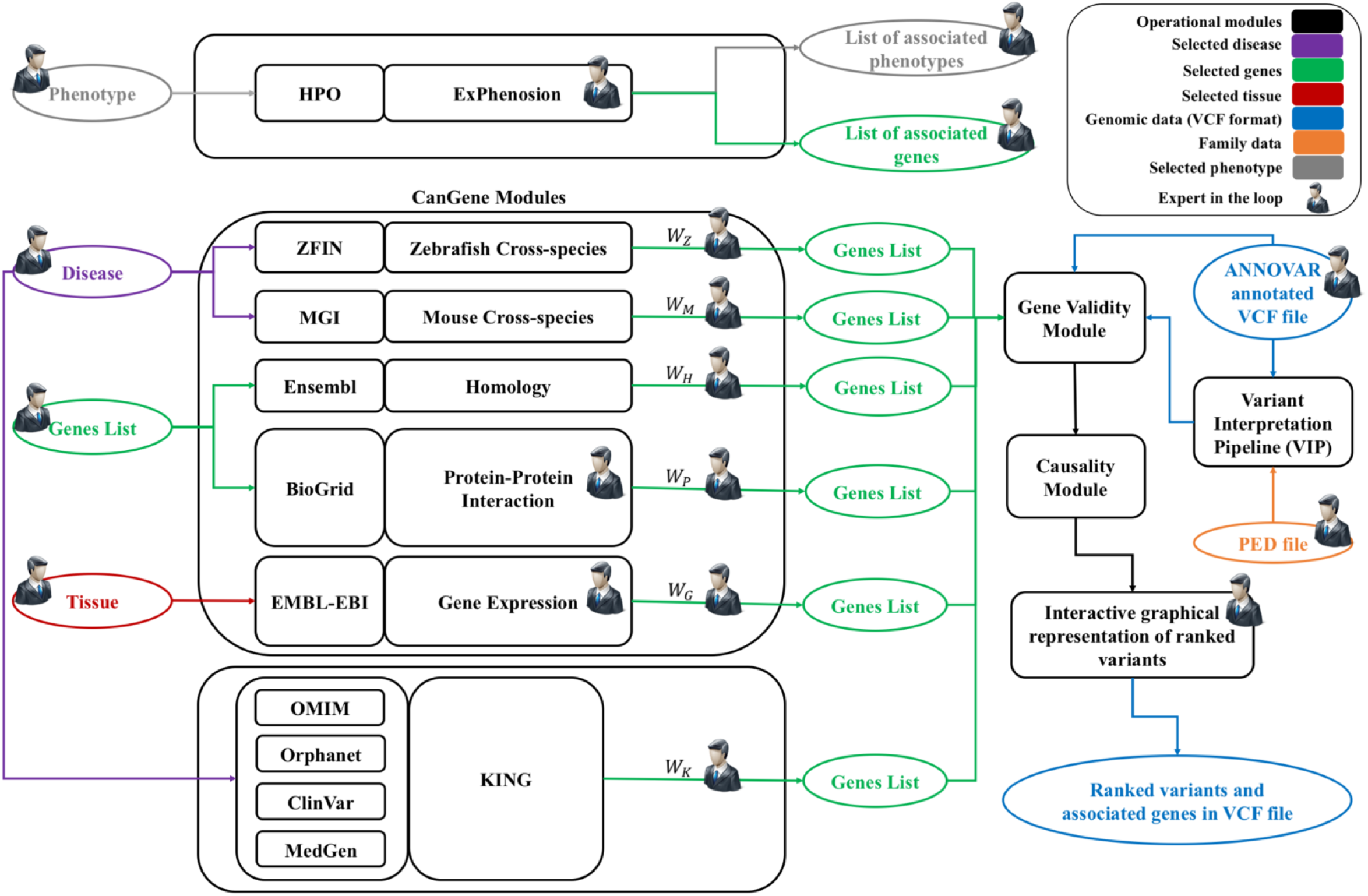
*GeneTerpret* workflow. This figure depicts the modules, their feeding databases, acceptable inputs, and the flow of information in the workflow with the main modules feeding to the gene validity, *VIP*, and causality modules. Three sets of modules are available within *GeneTerpret* for gene validity exploration; (1) ExPhenosion module - accepts the phenotype as an input; the number of super-classes to walk up can be customized, and the outputs return the connected phenotypes and their associated genes. This module works independently from other developed modules to extract the connected phenotypes to the selected phenotype and allow the analyst to explore the genes associated with related phenotypes; (2) CanGene modules - generate a list of candidate genes by compiling various types of evidence. The cross-species (Zebrafish and mouse) modules accept the disease(s) by its/their MONDO ID(s) as the input(s) and generate a list of genes that their orthologue is associated with similar disease in animal models by checking the related databases. The Homology and Protein-Protein Interaction modules accept a list of known genes for a phenotype (if it is available); the Homology module returns the homologous genes (paralogues) to the genes in the known gene list. The Protein-Protein interaction module takes a similar approach to generate a list of genes that interact with the known disease genes. The analyst can select the number of interaction neighborhood levels (such as level-1, level-2, etc.) desired for this interpretation. The Gene Expression module accepts a list of relevant tissues as input and outputs the list of genes expressed in the selected tissue based on the expression cut-off threshold which is set by the analyst; (3) KING module - accepts a disease(s) (MONDO ID(s)) as the input and then outputs a list of genes associated with the said disease based on evidence obtained from Orphanet, OMIM, ClinVar, and MedGen databases. The validity module accepts the generated gene lists from the modules CanGene and KING, as well as ANNOVAR, annotated VCF file or the output of VIP module as an input. The output file is the VCF file including validity scores or the VCF file including validity scores. VIP module has been developed based on ACMG guidelines.^16^ This module annotates the variants with pathogenicity terms (PVS1, PS1, etc.) and justifies the assigned terms. The causality module integrates the output of validity and VIP modules and ranks the variants based on the number of evidence extracted from validity modules and pathogenicity terms from VIP. Simultaneously, an interactive graphical representation of the variants is generated which allows the analyst to select the desired variants by using a LASSO filter.

### GeneTerpret modules and functions

#### A. Generating and Querying Known and Candidate Gene lists and Exploring Phenotype Associations

To establish gene-disease validity in *GeneTerpret*, the general interpretation workflow consists of three modules that extract the phenotype terms, their associated genes, and their candidate genes. The first module, Known INvolved Genes (*KING*), outputs a list of genes associated with a particular phenotype(s), with strong evidence of support from *OMIM*^11^, *Orphanet*^12^, *MedGen*^13^, and *ClinVar*^14^. The second module, Expanded Phenotype Exploration (ExPhenosion), uses the Human Phenotype Ontology (*HPO*) hierarchy of phenotype^15^ to produce a list of genes associated with a particular phenotype, (e.g. *Tetralogy of Fallot*), including superclass terms which is the broader phenotype for a specific HPO term (e.g. *Conotruncal defect*). The third module, Candidate Genes (*CanGene*), produces a list of candidate genes for a given phenotype by collecting various pieces of biological evidence from many relevant databases (Table 1). The details of each module and database used by *GeneTerpret* are described in the supplementary file (S2 section) and Supplementary Table S1.

**Table 1).** Databases used by operational modules in the *GeneTerpret* Platform.

| Module |  |  | Task | Databases |
| --- | --- | --- | --- | --- |
| ExPhenosion |  |  | Identifies genes associated with the selected phenotype(s) and its/their superclass phenotypes | Human Phenotype Ontology (HPO) <sup>15</sup><br>Medical Subject Headings (MeSH) <sup>24</sup> |
| CanGene | Cross-species | Mouse | Identifies candidate genes that cause a “similar” phenotype in a mouse model | Mouse Genome Informatics (MGI) <sup>25</sup> |
|  |  | Zebrafish | Identifies candidate genes that cause a “similar” phenotype in the zebrafish model | The Monarch Initiative<sup>26</sup> |
|  | Homology |  | Identifies candidate genes homologous to known disease genes for a phenotype | Ensembl<sup>27</sup> |
|  | Protein-Protein interaction |  | Identifies candidate genes that physically interact with known disease genes in human studies | The Biological General Repository for Interaction Datasets (BioGRID) <sup>28</sup> |
|  | Gene Expression |  | Identifies candidate genes expressed in the affected tissue | EMBL-EBI Expression Atlas <sup>29</sup> |
| Known INvolved Genes (KING) |  |  | Identifies known genes for a selected phenotype | Online Mendelian Inheritance in Man (OMIM) <sup>11</sup><br>Orphanet <sup>12</sup><br>NCBI MedGen <sup>13</sup><br>NCBI ClinVar <sup>14</sup> |
| Gene Validity |  |  | Calculates validity scores for each gene by examining the strength of the evidence supporting a gene-disease relationship obtained from the above modules | N/A |
| Variant Interpretation Program (VIP) |  |  | Classifies variants based on their pathogenicity following the criteria proposed by the American College of Medical Geneticists (ACMG) | ClinGen Dosage Sensitivity Map <sup>30</sup><br>Decipher haploinsufficiency predictions <sup>31</sup><br>ExAC pLI score <sup>32</sup><br>ClinVar <sup>14</sup><br>The NHGRI-EBI Catalog of published GWAS <sup>33</sup><br>Pfam clans <sup>34</sup><br>Laurens W et al, 2017 <sup>35</sup> |
| Causality |  |  | Graphical visualization of the distribution of prioritized variants across the five classifications of pathogenicity | GeneTerpret GUI |

#### B. Gene Validity Module - Integration of Validity Terms

The gene validity module is used to quantify the strength of evidence that supports a gene-disease relationship. This module consolidates output gene lists from *CanGene, ExPhenosion*, and *KING* modules (see gene validity module architecture in Supplementary Figure S2), and appends a score for each gene based on the number of times it appears in the output of the modules, the user-assigned weights for each module, and the user-defined thresholds. We recommend that *KING* output be considered as strong evidence (known genes) as it is based on published genes associated human phenotypes in the four common medical genetics databases, while the other outputs can be treated as limited evidence (candidate gene). The acceptable inputs for gene validity modules are (a) gene lists obtained from any of *CanGene*, or *KING* module, (b) uploaded gene-list, or (c) uploaded VCF file. The module output provides a new, annotated VCF file with added column(s) showing the weight of evidence for each gene from the list previously generated from each selected module. For each gene, a validity score that summarizes all evidence is also given in the output. It is important to note that the module parameters (such as the thresholds and weights) set by the analyst will impact the validity scores produced.

#### C. Variant Interpretation Program (*VIP*) Module - Determining Variant Pathogenicity

The Variant Interpretation Program (*VIP*) module establishes and appends pathogenicity calls to variants from a given VCF file. The internal structure of this module has been shown in Supplementary Figure S3, and the respective databases used in this module are presented in the Supplementary Table S1. This module accepts a VCF file in *ANNOVAR* annotation format (Supplementary Table S2), and where family history is available, a combination of PED and VCF files as its input to achieve trio analysis (example in Supplementary Table S3). This module then outputs a set of new annotated VCF files, each with new columns added, showing ACMG pathogenicity classification for each variant, the ACMG criteria invoked, and the justification for arriving at a given classification. Overall, for each sample/trio/cohort analyzed, three VCF files are created; the first contains only de novo variants (if the family history is provided), the second lists only pathogenic and likely pathogenic variants, and the third lists all variants with pathogenicity classification. It is important to emphasize that variant pathogenicity classifications from *VIP* do not intend to conclusively indicate a variant’s clinical significance. The variant classifications from *VIP* are merely algorithmic predictions of pathogenicity based on the applications of the ACMG guidelines to each variant and hence does not mean the variant in question is conclusively pathogenic in a particular patient for the phenotype under consideration. Further details of the considered ACMG guidelines and their implementation can be found in Supplementary Table S4 of the 2015 ACMG publication.^16^

#### D. Causality Module – Visualization of The Interpreted Genomic Variants

Where there are many prioritized variants outputted from the *GeneTerpret VIP* module, we realized that a tool for the proper visualization of these variants would be helpful. Therefore, we developed the causality module which uses the output of the *VIP* and organizes the variants into a graphical visualization of the variants across the predicted pathogenicity categories plotted against the clinical validity scores of pertinent genes. This module is particularly helpful for visualizing prioritized variants when a high yield of prioritized variants is obtained. Supplementary Figure S4 shows a typical graph generated by the causality module which plots the variant distribution in the validity vs. pathogenicity space (Supplementary Figure S4(A)). The analyst can further filter the desired variants in this space by using an in-built lasso filtering tool (Supplementary Figure S4(B)).

### GeneTerpret Performance Assessment

To assess the performance of *GeneTerpret*, we did a performance assessment in two ways. First, we used two well-established external resources (ClinGen database^17,18^ for testing clinical validity modules and DECIPHER database^19^ for testing the variant pathogenicity module independently), and secondly, our expert-interpreted internal datasets composed on a Tetralogy of Fallot (TOF) cohort^20^ and Cardiac Genome Clinic (CGC) families^21^.

## RESULTS

We developed the *GeneTerpret* platform as a bioinformatics tool to facilitate the process of identifying disease-causing variants. Two orthogonal key concepts drive the interpretation of each variant: gene-disease clinical validity, and variant pathogenicity. Gene-disease validity is a qualitative measure of the strength of the evidence supporting the gene-disease relationship, quantifiable according to the ClinGen Gene Curation Project scale^22^ as no evidence, limited, moderate, strong, or definitive. For example, one can say SCN5A is “definitively” involved in “Brugada syndrome”, and that a high level of evidence supports the SCN5A-Brugada syndrome relationship.^23^ However, we designed *GeneTerpret* not to limit the user to these five categories; instead the platform allows the user to adjust the weight assigned to each source of evidence to produce a personalized validity factor based on the user’s preferences. Therefore, the variant pathogenicity outputted by the platform is a measure of the likelihood of a variant being detrimental to the gene/protein function. Pathogenicity in a clinical setting is expressed on a five-tier classification scale proposed by the ACMG: pathogenic, likely pathogenic, uncertain significance, likely benign, or benign. The causality is defined as the likelihood of a variant explaining the phenotype/disease observed in a patient. So, in *GeneTerpret*, a variant was considered “causal” when it ranked high on both gene-disease validity and variant pathogenicity scales. The details of datasets used and the design of the *GeneTerpret* package are described under methods.

### Validation of GeneTerpret’s Performance on External Data

#### A) Gene-Disease Validity Module

To validate the performance of the gene-disease validity module, we benchmarked it against the Gene-Disease clinical validity results from ClinGen. These are well established gene-disease associations curated by groups of experts in each field. Of 1082 curation records in the ClinGen Gene Validity curation table (https://search.clinicalgenome.org/kb/gene-validity accessed September 8, 2020), 715 were classified as “Definitive”, “Strong” or “Moderate” in association with 451 diseases. Running *GeneTerpret*’s KING module on these 451 diseases reproduced a list that contained 695 out of 715 (97.2%) genes in the ClinGen gene-validity table.

#### B) Performance of VIP

To benchmark the performance of *VIP*, we analyzed the entire DECIPHER dataset and compared the pathogenic/likely pathogenic and benign/likely benign annotations from DECIPHER with results obtained from *VIP*. A summary of the results of *VIP* for all the variants obtained from DECIPHER (8610 variants) is in Table 2. Interestingly, the percentage of variants called to be of uncertain significance were increased (42% in *VIP* vs. 38.6% in DECIPHER) at respectively a lower percentage of benign/likely benign calls (1.1% vs 2.7%). The percentage of pathogenic and likely pathogenic calls is relatively stable. Overall, there is high concordance (83.5%) between pathogenic or likely pathogenic calls classified by *VIP* and obtained by DECIPHER.

**Table 2).** *VIP* Interpretation of all variants from DECIPHER.

| **Clinical Significance** | **VIP (Automated Pathogenicity Identifier module)** | **DECIPHER (Manual Pathogenicity Identifier)** | **Concordant** |
| --- | --- | --- | --- |
| Benign | 14 (0.1%) | 23 (0.3%) | 0 (0%) |
| Likely Benign | 81 (0.9%) | 211 (2.4%) | 9 (0.2%) |
| Uncertain significance | 3633 (42.2%) | 3329 (38.6%) | 2202 (48.9%) |
| Likely pathogenic | 2692 (31.3%) | 2508 (29.1%) | 1055 (23.4%) |
| Pathogenic | 2190 (25.4%) | 2539 (29.5%) | 1240 (27.5%) |
| Sum of five tiers | 8610 | 8610 | 4506 |
| Benign or likely benign | 95 (1.1%) | 234 (2.7%) | 9 (0.2%) |
| Pathogenic or likely pathogenic | 4882 (56.7%) | 5047 (58.6%) | 3764 (83.5%) |

### *Validation of GeneTerpret*’s *Performance on Internal Data (Manually Prioritized Variants)*

To compare the performance of *GeneTerpret’s* variant classification with manual classifications by experienced genome analysts, blinded reinterpretations of two sample datasets were done using the *GeneTerpret* platform. The first set consisted of 10 families, and the second one consisted of 20 individuals, all from our internal database. To rank the *GeneTerpret*’s output, we employed a binning system based on scores from the gene-validity module and pathogenicity tier from *VIP* and sorted the variants into four bins. The bins were composed of; 1) pathogenic/likely pathogenic variant in a known (high validity) gene (P/LP KG); 2) pathogenic/likely pathogenic variant in a candidate (moderately valid) gene (P/LP CG); 3) pathogenic/likely pathogenic variant in a novel gene (P/LP NG); and 4) the variant of uncertain significance in a known gene (VUS KG) (Supplementary Table S5). Variants in each bin were further ranked based on the validity score of the corresponding genes. Results of validity scores from *GeneTerpret* were then compared with previous interpretations by our experienced geneticists; the latter findings were peer-reviewed and published already.^20^

#### A) GeneTerpret’s Performance in Family Interpretation

We tested 10 parent-child trios (VCF files) from a previous whole-genome sequencing (WGS) study of pediatric patients with cardiac phenotypes.^21^ In 4 of 10 families, the variant of interest (VOI) identified through manual curation was ranked among the top 10 variants in *GeneTerpret*’s output. Expanding the list to the top 50 ranked variants led to the inclusion of 9 out of the 10 final calls by an expert geneticist interpretation. Interestingly, *GeneTerpret* correctly identified all de novo variants from the families tested. Overall, 8 out of 12 *VIP* classified pathogenic variants were in complete agreement with results from previous manual interpretation. Three of the variants not in concordance with the previous expert-review included a variant each in *NIPBL, PTEN*, and *MYH11* gene from family FAM32, FAM13, and FAM54 respectively. The other discordance variant also in FAM34 (previously classified as likely pathogenic by manual interpretation) was re-classified a VUS by *VIP*: FLT4 (NM_182925.4) c.89delC, p.(Pro30Argfs*3) - frameshift variant did not fulfill the PM2 category of being rare/absent in controls (it had a minor allele frequency (MAF) of 5E-04 in gnomAD versus our stringently defined cut-off of MAF<1E-5). Figure 2(A) summarizes *GeneTerpret*’s output in comparison with the previously interpreted variants. Diseases/phenotypes used as the input of gene-validity modules to generate gene validity scores for each family are listed in Supplementary Table S6.

**Figure 2).**
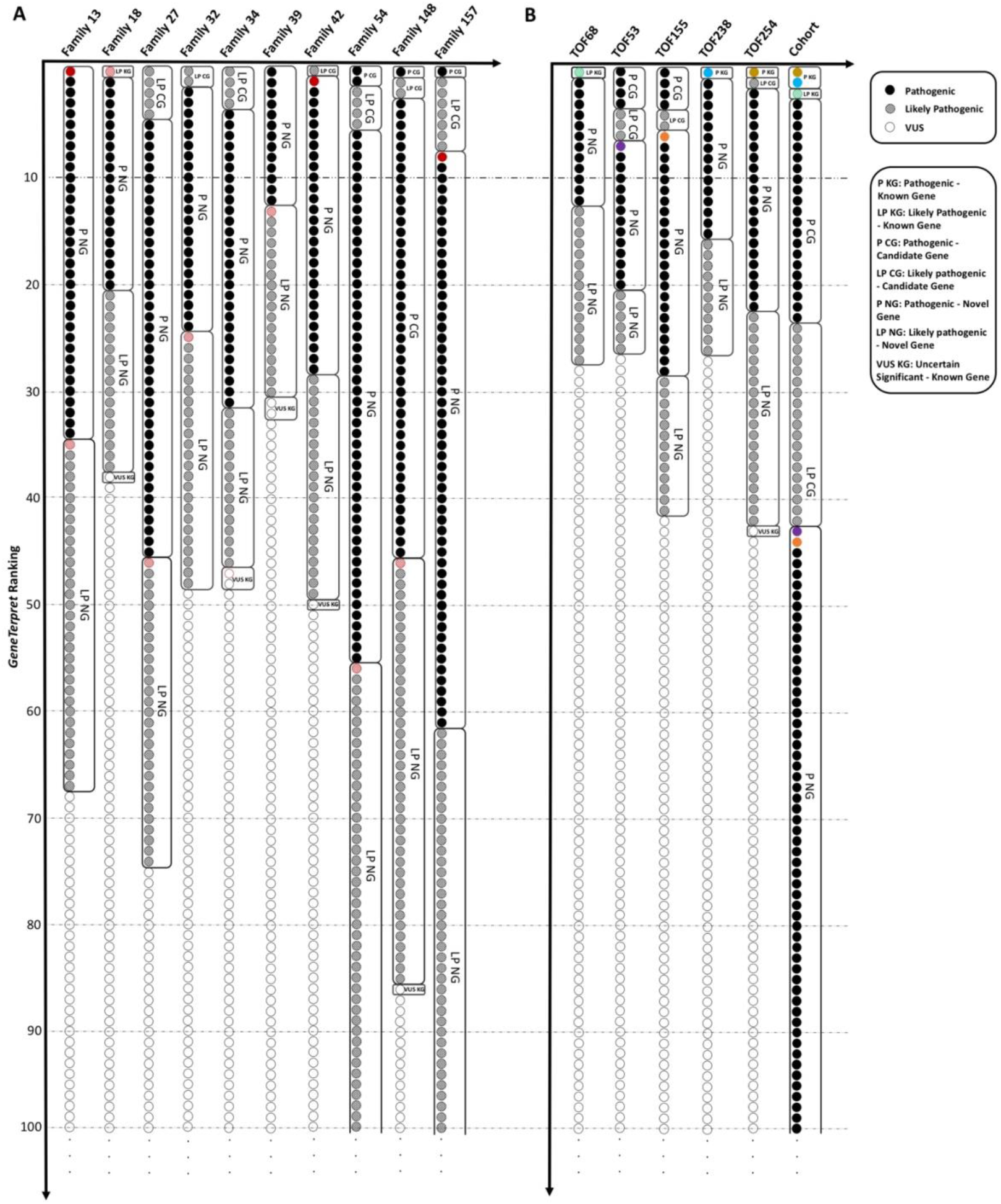
Graphical representation of the results from an analysis of internal datasets by *GeneTerpret* and manual interpretation. **(A)** The top hundred of ranked variants from the family-based analysis of ten families are represented. The red colour is highlighting the variant of interest (VOI) selected by a human analyst as published before^21^. The boxes around the variants cluster the same ranked variants by *GeneTerpret* (the same pathogenicity and validity terms). **(B)** The cohort-based results for 20 unrelated probands with “Tetralogy of Fallot”. The top hundred ranked variants are plotted as circles from top to bottom. The only five VOIs selected by a human genome analyst in five patients from this cohort^20^ are highlighted in colours. Different colours have been selected to distinguish the VOI related to each patient. For comparison, individual analysis of genomes from the five probands with VOIs are also plotted using the same colour-coding. For instance, the purple colour represents the obtained VOI for patient TOF53 (one of the probands in the cohort). This variant is ranked 44 in the cohort-based analysis and ranked 8 in the singleton-based analysis by *GeneTerpret*.

#### B) GeneTerpret’s Interpretation Performance in a Cohort of Individual Samples

To assess *GeneTerpret’s* ability to process multiple VCF files from a cohort of individual samples, we analyzed a dataset containing 20 unrelated probands, including five that had VOI findings as published in a previous study of tetralogy of Fallot.^20^ Figure 2(B) and Supplementary Table S7 summarize the results obtained from both the cohort-based and individual analysis. Interestingly, there was a complete agreement between *GeneTerpret’s* based-classification with expert geneticist’s manual interpretation. Moreover, the VOI was always in the top 50 variants in *GeneTerpret’s* output (three out of five ranked within the top 10 ranked variants). *GeneTerpret* also classified additional variants as pathogenic, likely pathogenic in candidate and novel genes, as well as variants of unknown significance in candidate genes. Important to note that the clinical significance of these additional variants requires a review by an analyst and/or further lab investigation. They may contain potential secondary findings, additional causal variants or modifiers for the phenotype of interest.

## DISCUSSION

*GeneTerpret* is a visual analytics platform that utilizes information from a variety of databases and modules to assist in speeding up the laborious process of genome variant interpretation (GVI). This platform was designed and implemented to streamline and optimize the expert genome analysis process by automating many of the data gathering, comparison, and filtration steps in the GVI process. To computationally achieve this, we re-packaged the data and computational tools into workable, and tunable modules that can be connected to different pipeline networks through an intuitive graphical user interface (GUI). This GUI also allows the user to tailor the platform output by adjusting the connections between modules within the library to suit their needs. Although *GeneTerpret* helps to automate much of the genome interpretation process, the user analysts remain key, especially in exploring candidate genes relevant to the desired phenotype and tissue, adjusting the weight of each validity term, and performing the final ranked output review. The investigative process of connecting the pieces of evidence among seemingly disconnected modules reflects various strategies that different genome analysts employ to decipher the causal genes and organize the genomic variants based on phenotypic information.

*GeneTerpret* is encouragingly accurate when compared with expert-curated datasets in such well-established public records of clinically relevant variants as DECIPHER and ClinGen. For the pathogenic or likely pathogenic variants in DECIPHER, *GeneTerpret*’s *VIP* showed a high concordance (83.5%) in calling variant pathogenicity. When it comes to gene-disease validity, the KING module showed an extremely high agreement (97.4%) with ClinGen expert-curated table to pull out genes with moderate to strong evidence for 451 diseases. Therefore, *GeneTerpret* should significantly facilitate the clinical genome interpretation process once publicized and adopted by genome analysis and interpretation labs across the world. Based on the analysis of our internal and equally expert-reviewed data, the variants of interest were mostly ranked in the top 10 *GeneTerpret*’s output (top 50 for all cases except one). These results highlighted the ability of *GeneTerpret* to efficiently analyze singletons, trios or cohorts by generating a manageable, prioritized list of variants for further in-depth interpretation based on the phenotype, essentially cutting down on typical analysis time from days or hours to minutes. By re-adjusting thresholds to include more stringent cut-offs, *GeneTerpret* ranking improves significantly, as exemplified by all above-referred variants being ranked within the top 50 gene list. Likewise, the results from cohort analysis demonstrated the ability of *GeneTerpret* to accelerate genome interpretation significantly. Indeed, *GeneTerpret* was able to analyze up to 30 samples within a few minutes and generate a robust, manageable ranked list of variants. However, we caution that using the *GeneTerpret* platform does not obviate the need for a human interpreter, but rather, it is a platform that provides an efficient aid to aggregate validity evidence and rank the variants, thus significantly reducing the time needed by an interpreter to sieve through an unsorted VCF file. *GeneTerpret* would significantly reduce the time required for the interpretation of large genomic datasets, particularly for large cohort analysis, which can take months to analyze.

Although a growing number of commercial or free packages are now in use to aid in classifying germline variants following the ACMG criteria, these tools are often too complex for routine clinical use, force the users to accept and follow the designed routine, and do not provide enough information for the users to draw meaningful conclusions from their predictions. These tools are often designed as black-box (closed-box) systems where the user is given minimal knowledge of the system architecture, so the users cannot gain access to the internal modules of the system. In fact, most of their key parameters and databases of the internal modules remain hidden or not freely available. Therefore, users of such commercial tools only learn or know the required inputs and expected outputs, with less information on how the tools or platforms function. We believe that an ideal genetic data analysis platform should be flexible, with tools designed as a white-box system for the users to see and interact with, and hence allowing for full interactive information flow. Additionally, it should allow the users to analyze the data and information flow, and control parameters and databases within the system. However, implementation of a white-box system is hard and would be computationally impossible on a web-based platform. So, to balance the accuracy, speed and efficiency of the system, we developed *GeneTerpret* as a gray-box system, which splits the difference between white-box and black-box systems and balances the user’s engagement time and the level of information. Our design attempts to not only allow users to understand the system and access the designed modules in the library but also to provide a workspace environment to check the result of each module independently while showing the users which modules can be connected to make their interpretation routine meaningful.

Over time, more user preferences and analytical options will be included as computational abilities and technologies continue to allow. So, we know that *GeneTerpret*, in its current version, has a few notable limitations. First, it is limited to analyzing single nucleotide variants (SNVs) with no functionality to analyze copy number or structural variations. Second, its functionality in analyzing familial data is limited to trios. Third, given the rigidity of some of the criteria (such as population frequency and haploinsufficiency cut-offs), the final call may differ from that conducted by an expert genetic variant interpreter (geneticist) who understands more nuanced scenarios. Finally, some of the parameters and filters for pathogenicity and validity are not customizable. We intend to provide more customization and interactive visual feedback in the future. Nonetheless, we believe that *GeneTerpret* will not only make the genome process streamlined, but will help facilitate the gene discovery process. Importantly, *GeneTerpret* more effectively addresses two main challenges: (1) it reduces the time of interpretation significantly by collecting evidence and sorting variants, and; (2) it provides a visual, flexible workspace for the analyst to develop and customize their routine.

## Supplementary Tables and Figures

**Supplementary Table S1) *GeneTerpret modules and respective databases with links to the used data***

**Supplementary Table S2) The required VCF file annotation, headers and descriptions**

**Supplementary Table S3) The standard format for the PED file**

**Supplementary Table S4) Variant Interpretation Program (*VIP*) logic (pseudocode) for variant classification following ACMG criteria**

**Supplementary Table S5) Bins Used for Ranking *GeneTerpret* Output**.

**Supplementary Table S6) Comparison of *GeneTerpret* output and previous manual interpretation of 10 trios**.

**Supplementary Table S7) Comparison of *GeneTerpret* output and previous manual interpretation of a cohort with 20 TOF patients**. Findings from manual interpretation have been reported for five individuals in this cohort.

**Supplementary Figure S1) A snapshot of the *GeneTerpret* graphical user interface (*GeneTerpret GUI*)**. A general interpretation routine is depicted as an example. The user selects the needed modules from the top right panel; then drags and drops them one by one in the left workspace panel. Furthermore, the tissue or phenotype/disease of interest can be directly entered by the user as an input in the bottom right panel and the generated module could be dragged and dropped in the left workspace panel. The users can upload their annotated VCF file, gene list(s), family information (PED file) and phenotypes/diseases list as further input for *GeneTerpret* by tapping on the upload tab in the bottom right panel and drag and drop the assigned generated module for the uploaded file in the workspace panel in the left side.

**Supplementary Figure S2) Gene Validity Module architecture**.

**Supplementary Figure S3) Variant Interpretation Program (*VIP*) internal structure**.

**Supplementary Figure S4) Overview of the causality module output**; **(A)** the interactive visualization of variant distribution in the validity-pathogenicity space allows users to explore the desired variants. Dark green, light green, yellow, orange, and red colours represent the pathogenicity of variants in a 5-tier system: benign, likely benign, uncertain significance, likely pathogenic, and pathogenic variants. **(B)** Lasso filter allows the analyst to select the desired variants and filter them to a downloadable VCF file.

## Data Availability

*GeneTerpret* is a web-based interpretation tool available online at https://geneterpret.com/

## Acknowledgements

We would like to thank our colleagues (specifically Dr. Miriam S. Reuter, Reem Khan, Cara Murphy, Dr. Anne S. Bassett, Dr. Daniele Merico, Dr. Christian R. Marshall, and Eriskay Liston) for their constructive remarks to improve the website and to provide including additional case studies. This study makes use of data generated by the DECIPHER community. A full list of centres who contributed to the generation of the data is available from https://decipher.sanger.ac.uk/about/stats and via email from. Funding for the DECIPHER project was provided by the Wellcome Trust foundation. This work was funded by generous support from the Ted Rogers Centre for Heart Research at The Hospital for Sick Children.

## Author Information

RM and SMH designed this framework. RM led the development process and analysis. RM, SMH led the analysis interpretation and manuscript writing. SD and VA contributed extensively to the development process. PD contributed to the development process. SD and EJ contributed extensively to analysis interpretation and manuscript writing. EJ designed the website. JBAO, KK, KMF, CS, RKJ, and SWS contributed to the analysis, interpretation and revision of the manuscript. RHK and SMH coordinated the project and provided overall leadership. All authors discussed the results, provided critical feedback, and contributed to the final manuscript, and approved the submitted version.

## Ethics Declaration

Internal data used in the validation part of this study was obtained as part of research protocols approved by the Research Ethics Boards at The Hospital for Sick Children (REB #1000053844), the University Health Network (REB 98-E156), and the Centre for Addiction and Mental Health (REB 154/2002).

