## Supplementary information for "GeneTerpret: a customizable multilayer approach to genomic variant prioritization and interpretation"

### Supplementary Materials

#### S1. *GeneTerpret* Backend and Web Implementation

*GeneTerpret GUI* has been implemented in HTML5 and JavaScript, using Paper.js 0.11.5[1], jQuery 3.2.1[2], Bootstrap 3.3.7[3], and Plotly 1.49.4[4] libraries. As nodes are created, connected, modified or tuned (via parameters), *GeneTerpret* sends HTTP requests to the associated algorithms/tools in the backend. In turn, this determines if the processing is needed as directed by the connected nodes, and where criteria for processing are met, the platform then returns data to any nodes that have successfully executed. Multiple queries can be executed at the same time by connecting different nodes, hence allowing for parallel processing of independent modules.

*GeneTerpret*'s web server backend is coded in Python 3.7[5] using Flask 0.12[6] library. The web server communicates with a co-located MongoDB 3.4.9[7] server, which is used to store all user session data and the latest versions of external databases needed for processing. *MongoDB* was chosen over a relational database such as MySQL because the format of data across all sources varies greatly, and largely consists of nested or aggregated relationships. Flask library implements the Python Web Server Gateway Interface (WSGI) so that production-ready HTTP servers can expose the modules' processing on a large scale. In practice, we deployed the *GeneTerpret* backend to a Windows 2016 Server edition Virtual Private Server (VPS), hosted by OVHCloud[8]. The VPS has two 3 GHz virtual processing cores, 4 GB of RAM, and 50 GB of storage space.

#### S2. Details of implemented modules under validity module

##### *Module 1: ExPhenosion - Identification of Genes Associated with Superclass Phenotypes*

The *ExPhenosion* (Expanded Phenotype Exploration) module identifies the genes associated with a queried human phenotype(s) and expands the search by further exploration to cover broader phenotypes. The module uses data from the Human Phenotype Ontology (*HPO*), version 1.6.0 and Medical Subject Headings (MeSH) to generate a list of broader phenotypes (superclass) encompassing the original phenotype. For example, an input of "Tetralogy of Fallot" will return "Conotruncal defects" and "Heart Defects, Congenital" as the superclass. The module then returns a list of genes associated with the output phenotype(s) via a query of the *HPO* database. The analyst can modify the number of levels to search upstream of the input phenotype(s) in a hierarchy to adjust the breadth of the search.

##### *Module 2: KING - Identification of Known Involved Genes*

The Known INvolved Genes (*KING*) module finds known (high evidence) genes for a particular phenotype/disease. This module uses data from *OMIM*[9], *Orphanet*[10], *ClinVar*[11], and *MedGen*[12]. *OMIM* is an authoritative source of human genes and phenotypes/diseases that is updated daily. *Orphanet* provides genes related to a rare disorder to better describe rare disorders of genetic origin which are periodically updated. *ClinVar* is a genetic variant database and used as a resource to study genotype-phenotype correlations which are updated weekly. *MedGen* is a portal to information on genetic aspects of human health and disease which is updated daily. The output of *KING* is a gene list, which can be connected to the validity module as an input.

***Module 3: CanGene - Identification of Candidate Genes.***

*CanGene* assembles a list of candidate genes using four sub-modules based on protein-protein interactions, sequence homology, gene expression, and cross-species comparisons.

***I. Cross-Species modules***

The mouse and Zebrafish have many genetic similarities to humans. The protein-coding regions of mouse and human are on average 85% identical[13] and Zebrafish and human genomes are on average 70% identical[14]. Indeed, 84% of genes associated with human disease have a zebrafish counterpart[15]. Therefore, there are potentially interesting genes conserved across species that act in the same in these model organisms as would be in humans.

Our mouse and Zebrafish cross-species modules produce a list of candidate genes for a phenotype/disease(s) when the orthologues gene causes “similar” phenotypes in animal models (mouse or zebrafish). These modules query the MGI (v6.15), Monarch, and ZFIN databases[16].

***II. Homology module***

The Homology module provides a list of candidate genes that are homologues (sequence homology), to the known genes for a phenotype. The input is a list of known genes for the phenotype (user-provided or output of *KING*). This module uses a copy of the *ENSEMBL Paralogues* table to return other species genes that are homologous to the list of the imported genes.

***III. Protein-Protein Interactions module***

The Protein-Protein Interactions module provides a list of candidate genes that physically interact with the known genes for a phenotype. This module uses data from *BioGRID (3.5.181)* filtered for physical interactions and *Homo sapiens* to find all genes with direct interactions with the known genes. A user-provided list or a gene list obtained from other modules (*KING* in particular) could be used as the input. The user can adjust the number of publications that support each protein-protein interaction (default is 1).

***IV. Gene Expression module***

The Gene Expression module provides a list of candidate genes expressed in a particular tissue(s), relevant to the phenotype or disease of interest. This module uses experiment files fetched from the EMBL-*EBI Expression Atlas* database. The user can adjust the level of expression for a gene to get into the output list which could range from 1 (high expression, shorter list) to 3 (low expression, longer list).

**Supplementary Table S1)** *GeneTerpret* modules and respective databases and links/filters

| Module | Database Name | Database Source | Version Used |
|---|---|---|---|
| *ExPhenosion* | HPO | http://purl.obolibrary.org/obo/hp.obo | hp/releases/2020-02-27 |
| | HPO (phenotype to gene reference) | http://compbio.charite.de/jenkins/job/hpo.annotations/lastSuccessfulBuild/artifact/util/annotation/phenotype_to_genes.txt | Build #1270 |
| | MeSH | ftp://nlmpubs.nlm.nih.gov/online/mesh/2019/asciimesh/d2019.bin | d2019.bin |
| *CanGene: Mouse Cross-species* | Mouse Genomics Information | http://www.informatics.jax.org/downloads/reports/MGI_DO.rpt | 2020-03-09 09:04 |
| | Mouse Genomics Information (mouse to human phenotype reference) | http://www.informatics.jax.org/downloads/reports/HMD_HumanPhenotype.rpt | 2020-03-09 09:04 |
| *CanGene: Zebrafish Cross-species* | Monarch Initiative | https://solr.monarchinitiative.org/solr/golr/select/?defType=edismax&qt=standard&indent=on&wt=csv&rows=100000&start=0&fl=subject,subject_label,subject_taxon,subject_taxon_label,object,object_label,relation,relation_label,evidence,evidence_label,source,is_defined_by,qualifier&facet=true&facet.mincount=1&facet.sort=count&json.nl=arrarr&facet.limit=25&facet.method=enum&csv.encapsulator=%22&csv.separator=%09&csv.header=true&csv.mv.separator=%7C&fq=subject_category:%22gene%22&fq=object_closure:%22HP:0001626%22&fq=subject_taxon_label:%22Danio%20rerio%22&facet.field=subject_taxon_label&q=*:* | 2020-02 |
| | Monarch Initiative (zebrafish to human gene reference) | https://zfin.org/downloads/ortho.txt | 8 Mar 2020 |
| *CanGene: Homology* | Ensembl Paralogue table | http://www.ensembl.org/biomart/martservice?query=%3C?xml%20version=%221.0%22%20encoding=%22UTF-8%22?%3E%20%3C!DOCTYPE%20Query%3E%20%3CQuery%20virtualSchemaName%20=%20%22default%22%20formatter%20=%20%22TSV%22%20header%20=%20%221%22%20uniqueRows%20=%20%220%22%20count%20=%20%22%22%20datasetConfigVersion%20=%20%220.6%22%20%3E%20%3CDataset%20name%20=%20%22hsapiens_gene_ensembl%22%20interface%20=%20%22default%22%20%3E%20%3CAttribute%20name%20=%20%22ensembl_gene_id%22%20/%3E%20%3CAttribute%20name%20=%20%22external_gene_name%22%20/%3E%20%3CAttribute%20name%20=%20%22hsapiens_paralog_ensembl_gene%22%20/%3E%20%3CAttribute%20name%20=%20%22hsapiens_paralog_associated_gene_ | v0.6 |

| | | | |
|---|---|---|---|
| | | name%22%20/%3E%20%3CAttribute%20name%20=%20%22hsapiens_paralog_perc_id%22%20/%3E%20%3CAttribute%20name%20=%20%22hsapiens_paralog_perc_id_r1%22%20/%3E%20%3C/Dataset%3E%20%3C/Query%3E | |
| *CanGene: Protein-protein Interaction* | The BioGrid | https://downloads.thebiogrid.org/Download/BioGRID/Latest-Release/BIOGRID-ORGANISM-LATEST.tab2.zip | BIOGRID-3.5.182 |
| *CanGene: Gene Expression* | Ebi Expression Atlas (experiment MTAB 2706) | https://www.ebi.ac.uk/gxa/experiments-content/E-MTAB-2706/resources/ExperimentDownloadSupplier.RnaSeqBaseline/tpms.tsv | Static link |
| | Ebi Expression Atlas (experiment MTAB 2836) | https://www.ebi.ac.uk/gxa/experiments-content/E-MTAB-2836/resources/ExperimentDownloadSupplier.RnaSeqBaseline/tpms.tsv | Static link |
| | Ebi Expression Atlas (experiment MTAB 3358) | https://www.ebi.ac.uk/gxa/experiments-content/E-MTAB-3358/resources/ExperimentDownloadSupplier.RnaSeqBaseline/tpms.tsv | Static link |
| | Ebi Expression Atlas (experiment MTAB 3716) | https://www.ebi.ac.uk/gxa/experiments-content/E-MTAB-3716/resources/ExperimentDownloadSupplier.RnaSeqBaseline/tpms.tsv | Static link |
| | Ebi Expression Atlas (experiment MTAB 3871) | https://www.ebi.ac.uk/gxa/experiments-content/E-MTAB-3871/resources/ExperimentDownloadSupplier.RnaSeqBaseline/tpms.tsv | Static link |
| | Ebi Expression Atlas (experiment MTAB 4344) | https://www.ebi.ac.uk/gxa/experiments-content/E-MTAB-4344/resources/ExperimentDownloadSupplier.RnaSeqBaseline/tpms.tsv | Static link |
| | Ebi Expression Atlas (experiment MTAB 4840) | https://www.ebi.ac.uk/gxa/experiments-content/E-MTAB-4840/resources/ExperimentDownloadSupplier.RnaSeqBaseline/tpms.tsv | Static link |
| | Ebi Expression Atlas (experiment MTAB 513) | https://www.ebi.ac.uk/gxa/experiments-content/E-MTAB-513/resources/ExperimentDownloadSupplier.RnaSeqBaseline/tpms.tsv | Static link |
| | Ebi Expression Atlas (experiment MTAB 5214) | https://www.ebi.ac.uk/gxa/experiments-content/E-MTAB-5214/resources/ExperimentDownloadSupplier.RnaSeqBaseline/tpms.tsv | Static link |
| | Ebi Expression Atlas (experiment MTAB 5423) | https://www.ebi.ac.uk/gxa/experiments-content/E-MTAB-5423/resources/ExperimentDownloadSupplier.RnaSeqBaseline/tpms.tsv | Static link |
| | Ebi Expression Atlas (experiment PROT 1) | https://www.ebi.ac.uk/gxa/experiments-content/E-PROT-1/resources/ExperimentDownloadSupplier.Proteomics/tsv | Static link |
| | Ebi Expression Atlas (experiment PROT 3) | https://www.ebi.ac.uk/gxa/experiments-content/E-PROT- | Static link |

| | | | |
|---|---|---|---|
| | | 3/resources/ExperimentDownloadSupplier.Proteomics/tsv | |
| *Known INvolved Genes (KING)* | OMIM | http://omim.org/static/omim/data/mim2gene.txt | 2020-03-08 |
| | Orphanet | http://www.orphadata.org/data/xml/en_product6.xml | 01 Mar 20 |
| | MedGen | ftp://ftp.ncbi.nlm.nih.gov/pub/medgen/MedGen_HPO_OMIM_Mapping.txt.gz | 3/8/20 |
| | ClinVar | ftp://ftp.ncbi.nlm.nih.gov/pub/clinvar/tab_delimited/variant_summary.txt.gz | r2020-03-08 |
| *Variant Interpretation Program (VIP)* | ClinGen Dosage Map | https://ftp.clinicalgenome.org/ClinGen_haploinsufficiency_gene_GRCh37.bed | 2020-03-8 |
| | Decipher HI predictions | https://decipher.sanger.ac.uk/files/downloads/HI_Predictions_Version3.bed.gz | V3 |
| | Exac pLi | ftp://ftp.broadinstitute.org/pub/ExAC_release/release1/manuscript_data/forweb_cleaned_exac_r03_march16_z_data_pLI.txt.gz | r03_march16 |
| | ClinVar | ftp://ftp.ncbi.nlm.nih.gov/pub/clinvar/tab_delimited/variant_summary.txt.gz | 3/8/20 |
| | GWAS Catalogue | https://www.ebi.ac.uk/gwas/api/search/downloads/full | r2020-03-08 |
| | Pfam Clan List | ftp://ftp.ebi.ac.uk/pub/databases/Pfam/current_release/Pfam-A.clans.tsv.gz | 8/29/18 |
| | Journal PMC5656839 supplement | https://www.ncbi.nlm.nih.gov/pmc/articles/PMC5656839/bin/HUMU-38-1454-s003.xlsx | 2017 Aug 31 |

**Supplementary Table S2) The required VCF file annotation, headers and descriptions**

| annotations | Description |
|---|---|
| effect | type of effect on the coding sequence: (a) “synonymous SNV”, (b) "nonsynonymous SNV", (c) "stopgain SNV", (d) "frameshift deletion", (e) "frameshift insertion", (f) "frameshift substitution", (g) "nonframeshift deletion", (h) "nonframeshift insertion", (i) "nonframeshift substitution", (j) “stoploss SNV”. For variants with multiple effects, all possible values will be represented in comma-separated fashion. |
| typeseq | type of sequence overlapped with known genes/transcripts and their coding / noncoding status: (a) “exonic” represents coding exons, (b) “exonic: splicing” represents the beginning/end of coding exons which may also affect splicing, (c) “splicing” represents core splicing site (2 bp on the intron side of intron-exon and exon-intron junctions), (d) "ncRNA_exonic" represents exons of non-coding RNA genes, (e) "ncRNA_splicing" represents core splicing sites of non-coding RNA genes, (f) “UTR5” represents 5’ untranslated region, (g) “UTR3” represents 3’ untranslated region, (h) “upstream” represents 1kb upstream of TSS, (i) “downstream” represents 1kb downstream of TSS and (j) “intergenic” represents intergenic regions (beyond upstream/downstream threshold(1kb)). For variants with multiple sequence overlaps (e.g. exonic for one transcript and intronic for other), all possible typeseq values will be listed in semicolon-delimited format (e.g. exonic; intronic). |
| dbscSNV_ADA_SCORE | Splice site prediction scores from dbscSNV (https://www.ncbi.nlm.nih.gov/pubmed/26555599) |
| dbscSNV_RF_SCORE | Splice site prediction scores from dbscSNV (https://www.ncbi.nlm.nih.gov/pubmed/26555599) |
| refseq_id | combined Annovar output on coding sequence mapping and effect, composed of: (a) for coding exotic changes ("typeset" exonic): gene official |

| | |
|---|---|
| | symbol : RefSeq ID : position in the coding sequence : amino acid change , [idem]; (b) for core splice site changes (typeseq "exonic"): gene official symbol (RefSeq ID : exon number : coding sequence position and change , [idem]) |
| gene_symbol | official gene symbol. |
| dbsnp | exact match (position, allele) to dbSNP. |
| pfam_annovar | overlap with coding sequence matching to a PFAM protein domain; can help further assess protein impact, but not meant for hard-filter. |
| Repeat | Repeatmasker annotation from UCSC. |
| Exac_mis_z | Missense Z-score from ExAc. |
| sift_score | SIFT score for predicted protein impact, values <= 0.05 correspond to damaging (can be interpreted as a p-value); note that SIFT is based on amino acid conservation/substitution rates inferred from protein sequence alignments. |
| PROVEAN_score | Amino acid substitution or indel prediction score from Provean software (http://provean.jcvi.org/index.php). Values < -2.5 corresponds to a damaging variant. |
| polyphen_score | Polyphen2 scores for predicted protein impact, values >= 0.95 correspond to damaging; note that Polyphen2 is based on a set of sequence and structural attributes, combined in a machine classifier trained on a data-set of positive cases, and thus it partially is complementary and partially correlated to SIFT and MA. |
| ma_score | mutation assessor scores for predicted protein impact, values >= 2 correspond to damaging; note that MA is based on amino acid conservation/substitution rates inferred from protein sequence alignments, additionally modelling protein family groupings, thus can be regarded as an improved version of SIFT (improved performance has been shown particularly for somatic variants), although to this date SIFT remains more popular / broadly used. |
| mt_score | Mutationtaster prediction scores. |
| CADD_phred | PHRED-like c-score ranking from CADD. |
| phylopMam_avg | value array of PhyloP nucleotide-level conservation inferred from the Placental Mammal genome group; values >= 1 indicate moderate conservation, values >= 2.5 indicate strong conservation; note that PhyloP is based uniquely on nucleotide substitutions and does not factor structural variation. |
| phylopVert100_avg | The average value of PhyloP nucleotide-level conservation inferred from the 100 Vertebrate genome group; this field is suitable to hard filter, especially for missense variants and non-frameshift substitutions; values >= 1.5 indicate moderate conservation, values >= 4 indicate strong conservation; note that PhyloP is based uniquely on nucleotide substitutions and does not factor structural variation. |
| phastCons_placental | PhastCons score for the Placental Mammal genome group, based on UCSC track (calculated by HMM scan of PhyloP values to infer conserved status); this is useful to assess conservation at the *regional level* (rather than PhyloP's *nucleotide level*); this field can be used as evidence *suggestive* of the functional relevance of a sequence (especially if non-coding), and thus could be used to hard filter non-coding variation. |
| Clinvar_SIG | Overall ClinVar significance code; "pathogenic" is the code of interest for rare disorders. Clinical significance values for all the individual submissions (SCVs) aggregated for the RCV record in ClinVar (https://www.ncbi.nlm.nih.gov/clinvar/docs/clinsig/) |
| Clinvar_ReviewStatus | The level of review supporting the assertion of clinical significance(https://www.ncbi.nlm.nih.gov/clinvar/docs/details/#review_status) |
| X1000g_all | allele frequency in the full 1000 Genome data-set |

| | |
|---|---|
| ExAC_Freq | allele frequency in the full ExAC 65000 data set. |
| gnomAD_exome_ALL | allele frequency in the full Genome Aggregation Database exomes. |
| gnomAD_genome_ALL | allele frequency in the full Genome Aggregation Database whole genome sequences. |
| spx_dpsi | Splice site prediction score from SPIDEX. The change in percentage exon inclusion reported as the maximum across tissues (https://www.ncbi.nlm.nih.gov/pubmed/25525159) |
| gerp_wgs | whole-genome GERP++ RS scores greater than 2. |
| X1000g_eur | allele frequency in the caucasian-European sub-set of the 1000 Genome |
| X1000g_amr | allele frequency in the mixed-background Latin Americans (e.g. Mexicans, Puerto Ricans, Peruvians) sub-set of 1000 Genome |
| X1000g_eas | allele frequency in the east-Asian sub-set of the 1000 Genome |
| X1000g_afr | allele frequency in the black-African sub-set of 1000 Genome (note African here indicates the ethnic background, not the actual geographical location of the population sample) |
| X1000g_sas | allele frequency in the south-Asian sub-set of the 1000 Genome |
| ExAC_AFR | allele frequency in the African/African American sub-set of ExAC 65000. |
| ExAC_AMR | allele frequency in the Latino sub-set of ExAC 65000. |
| ExAC_EAS | allele frequency in the East Asian sub-set of ExAC 65000. |
| ExAC_FIN | allele frequency in the Finnish sub-set of ExAC 65000 |
| ExAC_NFE | allele frequency in the Non-Finnish European sub-set of ExAC 65000. |
| ExAC_OTH | allele frequency in the other populations of ExAC 65000. |
| ExAC_SAS | allele frequency in the South Asian sub-set of ExAC 65000. |
| gnomAD_exome_AFR | allele frequency in the African/African American sub-set of gnomAD exomes. |
| gnomAD_exome_AMR | allele frequency in the Latino sub-set of gnomAD exomes. |
| gnomAD_exome_ASJ | allele frequency in the Ashkenazi Jewish sub-set of gnomAD exomes. |
| gnomAD_exome_EAS | allele frequency in the East Asian sub-set of gnomAD exomes. |
| gnomAD_exome_FIN | allele frequency in the Finnish sub-set of gnomAD exomes. |
| gnomAD_exome_NFE | allele frequency in the Non-Finnish European sub-set of gnomAD exomes. |
| gnomAD_exome_OTH | allele frequency in the other populations of gnomAD exomes. |
| gnomAD_exome_SAS | allele frequency in the South Asian sub-set of gnomAD exomes. |
| gnomAD_genome_AFR | allele frequency in the African sub-set of gnomAD WGS. |
| gnomAD_genome_AMR | allele frequency in the Latino sub-set of gnomAD WGS. |
| gnomAD_genome_ASJ | allele frequency in the Ashkenazi Jewish sub-set of gnomAD WGS. |
| gnomAD_genome_EAS | allele frequency in the East Asian sub-set of gnomAD WGS. |
| gnomAD_genome_FIN | allele frequency in the Finnish sub-set of gnomAD WGS. |
| gnomAD_genome_NFE | allele frequency in the Non-Finnish European sub-set of gnomAD WGS. |
| gnomAD_genome_OTH | allele frequency in the other populations of gnomAD WGS. |

**Supplementary Table S3) The standard format for the PED file:**
PED is a text file showing family relationships. Here is an example of a trio with an affected proband and unaffected parents. The title line is unnecessary.

| **Family_id** | **Sample_id** | **Father_id** | **Mother_id** | **Sex** | **Phenotype** |
|---|---|---|---|---|---|
| HSC_001 | 6711234 | 9145028 | 7234560 | 2 | 2 |
| HSC_001 | 7234560 | 0 | 0 | 2 | 1 |

| | | | | | |
|---|---|---|---|---|---|
| HSC_001 | 9145028 | 0 | 0 | 1 | 1 |

**Supplementary Table S4)** Variant Interpretation Program (VIP) logic (pseudocode) for variant classification following ACMG criteria

| Classification | VIP (*GeneTerpret*) |
|---|---|
| PVS1 | If the column “ExonicFunc.refGene” contains “nonsense”, “stopgain”, “frameshift”, “frameshift insertion”, “frameshift deletion”, “frameshift substitution” and does not contain “nonframeshift” then set PVS1_t1 = 1<br><br>If the column “Func.refGene” contains “splicing” or “ncRNA_splicing” and the column “dbscSNV_ADA_Score” and/or “dbscSNV_RF_Score” is greater than 1, then set **PVS1_t1 = 1**<br><br>If the column “refseq_id” contains "p.Met1" then set **PVS1 = 1**<br><br>If the column “refseq_id” reflects a single or multiexon deletion, then set **PVS1_t1 = 1**<br><br>Using the ClinGen Dosage Map database determine haploinsufficiency score in (2,3). Using the Decipher determine the HI index less than 20%. Using the Exac pLi database determine EXAC_PLI greater than 0.8. Find the union among these gene lists and if the gene is in this result then set **PVS1_t2 = 1**<br><br>If both PVS1_t1 = 1 and PVS1_t2 = 1 then set **PVS1 = 1** |
| PS1 | If the column “Func.refGene” contains "Exonic” and the column “ExonicFunc.refGene” contains "Nonsynonomous" then set **PS1_t1 = 1**<br><br>Extract p.* in refseq_id column and cross-reference with the Clinvar database of pathogenic variants, if this gene is present in the Clinvar data then set **PS1_t2 = 1** |
| PS2 | Using the .ped file provided by the user check that the proband zygosity is equal to “ref-alt” and both parent zygosity are equal to “hom-ref” and is so then set **PS2 = 1** |
| PS3 | Set **PS3 = 0** |
| PS4 | Using the user-provided phenotype.<br><br>Use one level in the MeSH database (or the user-provided number of levels)<br><br>Query the GWAS catalogue for the selected phenotype and filter by a p-value less than 0.05 and Odd Ratio (OR)<br><br>Pull SNPs ids in the temp file to make a temporary dictionary<br><br>For each variant, if the “dbsnp” column is in the temp dictionary, then set **PS4 = 1** |
| PM1 | If the value of the **“**ExonicFunc.refGene” column is anything except “synonymous”, “NA” or “unknown” and the “Func.refGene” column is “Exonic”, then set **PM1_t1 = 1**<br><br>If the “pfam_annovar” column has a value then concatenate the values for the “Chr”, “Gene”, and “pfam_annovar” columns with underscores between each value and if the result is in the dictionary defined by us* and it has a value of 1 the set **PM1_t2 = 0**; otherwise, if it cannot be found in the dictionary then set **PM1_t2** = **1**<br><br>*Using Journal PMC5656839 supplement filter the “MAD_tolerance_score” column greater than 0.1 and convert the values to interpro id using the Pfam clan information |
| PM2 | Find max 31 columns (X1000g_all X1000g_eur X1000g_amr X1000g_eas X1000g_afr X1000g_sas ExAC_Freq ExAC_AFR ExAC_AMR ExAC_EAS ExAC_FIN ExAC_NFE |

<table>
<tr><td></td><td>ExAC_OTH ExAC_SAS gnomAD_exome_ALL gnomAD_exome_AFR gnomAD_exome_AMR gnomAD_exome_ASJ gnomAD_exome_EAS gnomAD_exome_FIN gnomAD_exome_NFE gnomAD_exome_OTH gnomAD_exome_SAS gnomAD_genome_ALL gnomAD_genome_AFR gnomAD_genome_AMR gnomAD_genome_ASJ gnomAD_genome_EAS gnomAD_genome_FIN gnomAD_genome_NFE gnomAD_genome_OTH)<br><br>If allele frequency is greater than max 31 columns, then set PM2 = 1; otherwise, set PM2 = 0</td></tr>
<tr><td>PM3</td><td>Set PM3 = 0</td></tr>
<tr><td>PM4</td><td>If the “ExonicFunc.refGene” column contains "nonframeshift insertion", "nonframeshift deletion", or "stoploss” and the “repeat” column is equal to “NA”, then set PM4 = 1</td></tr>
<tr><td>PM5</td><td>If the column “Func.refGene” contains "Exonic" and “splicing” and the “ExonicFunc.refGene” column contains "Nonsynonomous" then set PM5_t1 = 1<br><br>Extract p.* in refseq_id column<br><br>Use pathogenic variant dictionary from PS1.<br>If p.* is in the dictionary for the same gene, then set PS1_t2 = 1</td></tr>
<tr><td>PM6</td><td>Using the. ped file provided by the user checks that the proband zygosity is equal to “ref-alt” and both parent zygosity are equal to “hom-ref” and is so then set PM6 = 1</td></tr>
<tr><td>PP1</td><td>Set PP1 = 0</td></tr>
<tr><td>PP2</td><td>If the column “ExonicFunc.refGene” contains “nonsynony” and the column “Exac_mis_z” is greater than or equal to 3.09, then set PP2 = 1</td></tr>
<tr><td>PP3</td><td>Convert all scores to soft and hard categories (NA,0,1,2) based on this table:
<table>
<tr><th></th><th>Soft (1 point)</th><th>Hard (2 points)</th></tr>
<tr><td>sift_score</td><td><0.05</td><td><0.01</td></tr>
<tr><td>PROVEAN_score</td><td><-2.5</td><td><-4.1</td></tr>
<tr><td>polyphen_score</td><td>>0.15</td><td>>0.85</td></tr>
<tr><td>ma_score</td><td>>=1.9</td><td></td></tr>
<tr><td>mt_score</td><td>>=0.5</td><td></td></tr>
<tr><td>CADD_phred</td><td>>10</td><td>>20</td></tr>
<tr><td>phylopPMam_avg</td><td>1</td><td>2.5</td></tr>
<tr><td>phylopVert100_avg</td><td>1.5</td><td>4</td></tr>
<tr><td>phastCons_placental</td><td>NA</td><td>>500</td></tr>
</table>
If the mean is greater than or equal to 1, then set PP3 = 1</td></tr>
<tr><td>PP4</td><td>Set PP4 = 0</td></tr>
<tr><td>PP5</td><td>If the column “Clinvar_seq” contains “pathogenic” and the column “Clinvar_reviewstatus” contains “multiple submitters” and “no conflicts”, then set PP5 = 1</td></tr>
<tr><td>BA1</td><td>If the allele frequency is greater than 5% in Exome Sequencing Project, 1000 Genomes Project, or Exome Aggregation Consortium by checking the following.<br><br>Find max 31 cols (X1000g_all X1000g_eur X1000g_amr X1000g_eas X1000g_afr X1000g_sas ExAC_Freq ExAC_AFR ExAC_AMR ExAC_EAS ExAC_FIN ExAC_NFE ExAC_OTH ExAC_SAS gnomAD_exome_ALL gnomAD_exome_AFR gnomAD_exome_AMR gnomAD_exome_ASJ gnomAD_exome_EAS gnomAD_exome_FIN gnomAD_exome_NFE gnomAD_exome_OTH gnomAD_exome_SAS gnomAD_genome_ALL gnomAD_genome_AFR gnomAD_genome_AMR gnomAD_genome_ASJ gnomAD_genome_EAS gnomAD_genome_FIN gnomAD_genome_NFE gnomAD_genome_OTH)<br><br>If max 31 columns is greater than %5, then set BA1 = 1; otherwise, set BA1 = 0</td></tr>
</table>

<table>
<tr><td>BS1</td><td>Check if the allele frequency is greater than expected for the disorder by doing the following.<br><br>Find min 31 cols (X1000g_all X1000g_eur X1000g_amr X1000g_eas X1000g_afr X1000g_sas ExAC_Freq ExAC_AFR ExAC_AMR ExAC_EAS ExAC_FIN ExAC_NFE ExAC_OTH ExAC_SAS gnomAD_exome_ALL gnomAD_exome_AFR gnomAD_exome_AMR gnomAD_exome_ASJ gnomAD_exome_EAS gnomAD_exome_FIN gnomAD_exome_NFE gnomAD_exome_OTH gnomAD_exome_SAS gnomAD_genome_ALL gnomAD_genome_AFR gnomAD_genome_AMR gnomAD_genome_ASJ gnomAD_genome_EAS gnomAD_genome_FIN gnomAD_genome_NFE gnomAD_genome_OTH)<br><br>If min 31 columns are greater than 10 times the allele frequency from PM2, then set BS1 = 1, otherwise, set BS1 = 0</td></tr>
<tr><td>BS2</td><td>Set BS2 = 0</td></tr>
<tr><td>BS3</td><td>Set BS3 = 0</td></tr>
<tr><td>BS4</td><td>Set BS4 = 0</td></tr>
<tr><td>BP1</td><td>Set BP1 = 0</td></tr>
<tr><td>BP2</td><td>Set BP2 = 0</td></tr>
<tr><td>BP3</td><td>If the column “ExonicFunc.refGene” contains "nonframeshift insertion" or "nonframeshift deletion" and the “repeat” column does not contain “NA”, then set BP3 = 1</td></tr>
<tr><td>BP4</td><td>Convert all scores to soft and hard categories (NA,0,1,2) based on this table:
<table>
<tr><th></th><th>Soft (1 point)</th><th>Hard (2 points)</th></tr>
<tr><td>sift_score</td><td><0.05</td><td><0.01</td></tr>
<tr><td>PROVEAN_score</td><td><-2.5</td><td><-4.1</td></tr>
<tr><td>polyphen_score</td><td>>0.15</td><td>>0.85</td></tr>
<tr><td>ma_score</td><td>>=1.9</td><td></td></tr>
<tr><td>mt_score</td><td>>=0.5</td><td></td></tr>
<tr><td>CADD_phred</td><td>>10</td><td>>20</td></tr>
<tr><td>phylopPMam_avg</td><td>1</td><td>2.5</td></tr>
<tr><td>phylopVert100_avg</td><td>1.5</td><td>4</td></tr>
<tr><td>phastCons_placental</td><td>NA</td><td>>500</td></tr>
</table>
If the mean is less than 0.5, then set BP4 = 1</td></tr>
<tr><td>BP5</td><td>Set BP5 = 0</td></tr>
<tr><td>BP6</td><td>If the column “Clinvar_seq” contains “benign” and the column “Clinvar_reviewstatus” contains “multiple submitters” and “no conflicts”, then set BP6 = 1</td></tr>
<tr><td>BP7</td><td>If the column “Func.refGene” contains "Exonic" and does not contain “splicing” and the column “ExonicFunc.refGene” contains "synonymous", then set BP7_t1 = 1<br><br>If the value of the column “dbscSNV_RF_SCORE” is less than 0.6 and the value of the column “dbscSNV_ADA_SCORE” is less than 0.6 the value of the column “spx_dpsi” is greater than -10, then set BP7_t2 = 1<br><br>If the value of the column “phylopPMam_avg” is less than 1 and the value of the column “phylopVert100_avg” is less than 1.5 and the value of the column “gerp_wgs” contains “NA”, then set BP7_t3 = 1<br><br>If BP7_t1 and BP7_t2 and BP7_t3 are 1 then set BP7 = 1</td></tr>
</table>

**Supplementary Table S5) Bins Used for Ranking *GeneTerpret* Output.**

| Indicator | Bin | Validity score cut-off | Modules supporting gene-validity |
|---|---|---|---|
| P/LP KG | Pathogenic/Likely Pathogenic variant in a Known Gene | validity score >=5 | Strong evidence (*KING*) |
| P/LP CG | Pathogenic/Likely Pathogenic variant in a Candidate gene | 1 =< validity score < 5 | Medium evidence (*CanGene*) |
| P/LP NG | Pathogenic/Likely Pathogenic in a Novel Gene (no prior evidence) | validity score = 0 | None |
| VUS KG | Variants of Uncertain Significance in a Known Gene | validity score >=5 | Strong evidence (*KING*) |

Classification of variants into pathogenicity tiers (P/LP/VUS) reflects an algorithmic application of ACMG criteria and does not confer clinical significance or actual pathogenicity of a variant in a given case.

**Supplementary Table S6) Comparison of *GeneTerpret* output and previous manual interpretation of 10 trios.**

| Identifier [a] | Phenotype of proband | Variant of interest (VOI) obtained from manual curation | Position of VOI in *GeneTerpret* output | | # of P/LP KG [b] | # of P/LP CG [b] | # of VUS KG [b] | # of P/LP NG [b] |
|---|---|---|---|---|---|---|---|---|
| | | | Rank | Bin | | | | |
| FAM13 | VSD, axial hypotonia, hypoventilation, infantile spasms, macrosomia, macrocephaly, global developmental delay | **Pathogenic**: *PURA* (NM_005859.4) c.812_814delTCT,p.(Phe271del) | 1 | P/LP NG | 0 | 0 | 0 | 68 |
| | | **Pathogenic**: PTEN (NM_000314.4) c..95G>A, p.(Gly132Asp) | 36 | P/LP NG | | | | |
| FAM18 | Dysplastic aortic and pulmonary valve, pulmonary stenosis, aortic regurgitation | **Likely pathogenic**: *PTPN11* (NM_002834.4) c.209A>G, p.(Lys70Arg) | 1 | P/LP KG | 1 | 0 | 1 | 37 |
| | | **Variant of uncertain significance**: NOTCH1* (NM_017617.5) | 105 | VUS NG | | | | |

| | | | | | | | | |
|---|---|---|---|---|---|---|---|---|
| | | c.1934G>A, p.(Cys645Tyr) | | | | | | |
| FAM27 | Aortic stenosis, dysplastic aortic valve, microcephaly, hypotonia global developmental delay | **Likely pathogenic:** NEXMIF (KIAA2022) (NM_001008537.2) c.1502delG, p.(Gly501Valfs*4) | 47 | P/LP NG | 0 | 5 | 0 | 70 |
| FAM32 | AVSD, hypotonia, global developmental delay, borderline microcephaly | **Pathogenic:** NIPBL (NM_133433.3) c.771+1G>A, p.? | 27 | P/LP NG | 0 | 2 | 0 | 48 |
| FAM34 | TOF, MAPCA, pulmonary atresia, non-confluent hypoplastic pulmonary arteries, congenital lymphedema | **Likely Pathogenic:** FLT4 (NM_182925.4) c.89delC, p.(Pro30Argfs*3) | 48 | VUS CG | 0 | 4 | 2 | 43 |
| FAM39 | Aortic coarctation, BAV, macrocephaly, transverse arch hypoplasia, scoliosis | **De novo Likely Pathogenic:** NR2F2 (NM_021005.3) c.671T>A, p.(Val224Asp) | 14 | P/LP NG | 0 | 0 | 2 | 31 |
| FAM42 | AVSD, pulmonary vein stenosis, polyhydramnion, choroid plexus cyst, micrognathia | **Pathogenic:** ANKRD11 (NM_013275.5) c.5238_5239delGC, p.(Pro1747Argfs*49) | 2 | P/LP NG | 0 | 1 | 1 | 49 |
| FAM54 | PDA | **Pathogenic:** MYH11 (NM_002474.2) c.4578+1G>A, p.? | 57 | P/LP NG | 0 | 6 | 0 | 101 |
| FAM148 | Aortic coarctation, LAA, hypoplastic transverse arch, aortic stenosis, VSD, BAV, developmental delay, ptosis | **De Novo Likely Pathogenic:** KMT2D (NM_003482.3) c.15673C>T, p.(Arg5225Cys) | 47 | P/LP NG | 0 | 3 | 1 | 83 |
| FAM157 | HLHS, global developmental delay, hypotonia | **De Novo Pathogenic:** POGZ (NM_015100.3) c.3403delG, p.(Glu1135Argfs*3) | 9 | P/LP NG | 0 | 8 | 0 | 110 |

The green colour shows that the manual curation is completely matching with VIP results and the red colour represents the discrepancy.

* AVSD, atrioventricular septal defect; VSD, ventral septal defect; LAA, left aortic arch; TOF, tetralogy of Fallot; MAPCA, major aortopulmonary collateral arteries, PDA, patent ductus arteriosus; BAV, bicuspid aortic valve, HLHS, hypoplastic left heart syndrome.
[a] Identifiers of families described in Reuter *et al* 2020.
[b] Full list of additional variants and their classifications identified by *GeneTerpret* for these families are not reported because of secondary findings.

**Supplementary Table S7) Comparison of *GeneTerpret* output and previous manual interpretation of a cohort with 20 TOF patients.** Findings from manual interpretation have been reported for five individuals in this cohort.

| Identifier [a] | Disease/Phenotype (s) | Variant of interest (VOI) obtained from manual curation | Position of VOI in *GeneTerpret* output | | Count of variants [b] | | | |
|---|---|---|---|---|---|---|---|---|
| | | | Rank | Bin | P/LP KG | P/LP CG | VUS KG | P/LP NG |
| Cohort | TOF | FLT4 (NM_182925.4) c.1172_1173delAG, p.(Glu391Glyfs*35) | 1 | P/LP KG | 3 | 40 | 4 | 664 |
| | | FLT4 (NM_182925.4) c.2499C>G, p.(Tyr833*) | 1 | P/LP KG | | | | |
| | | KDR (NM_002253.2) c.2638C>T, p.(Arg880*) | 44 | P/LP NG | | | | |
| | | PRDM1 (NM_001198.3) c.1824C>A, p.(Cys608*) | 44 | P/LP NG | | | | |
| | | FLT4 (NM_182925.4) c.1037delC, p.(Thr346Argfs*7) | 3 | P/LP KG | | | | |
| TOF254 | TOF, APV; bilateral femoral vein occlusions; depression and/or anxiety | FLT4 (NM_182925.4) c.1172_1173delAG, p.(Glu391Glyfs*35) | 1 | P/LP KG | 1 | 1 | 1 | 41 |
| TOF238 | TOF, RAA, MAPCA, PA; aortic dilatation | FLT4 (NM_182925.4) c.2499C>G, p.(Tyr833*) | 1 | P/LP KG | 1 | 0 | 0 | 26 |
| TOF155 | TOF, PFO or ASD; depression and/or anxiety; gastroesophageal reflux | KDR (NM_002253.2) c.2638C>T, p.(Arg880*) | 7 | P/LP NG | 0 | 6 | 0 | 36 |
| TOF53 | TOF, PFO or ASD; coronary artery bypass grafts, systemic arterial hypertension; depression and/or anxiety | PRDM1 (NM_001198.3) c.1824C>A, p.(Cys608*) | 8 | P/LP NG | 0 | 7 | 0 | 20 |
| TOF68 | TOF, RAA, APV; depression and/or anxiety | FLT4 (NM_182925.4) c.1037delC, p.(Thr346Argfs*7) | 1 | P/LP KG | 1 | 0 | 0 | 27 |

All variants of interest were heterozygous. APV, absent pulmonary valve; ASD, atrial septal defect; PFO, patent foramen ovale; RAA, right aortic arch; TOF, tetralogy of Fallot; MAPCA, major aortopulmonary collateral arteries.

[a] Identifiers of individuals described in Reuter *et al* 2019.

[b] Count of variants in each bin of *GeneTerpret* output.

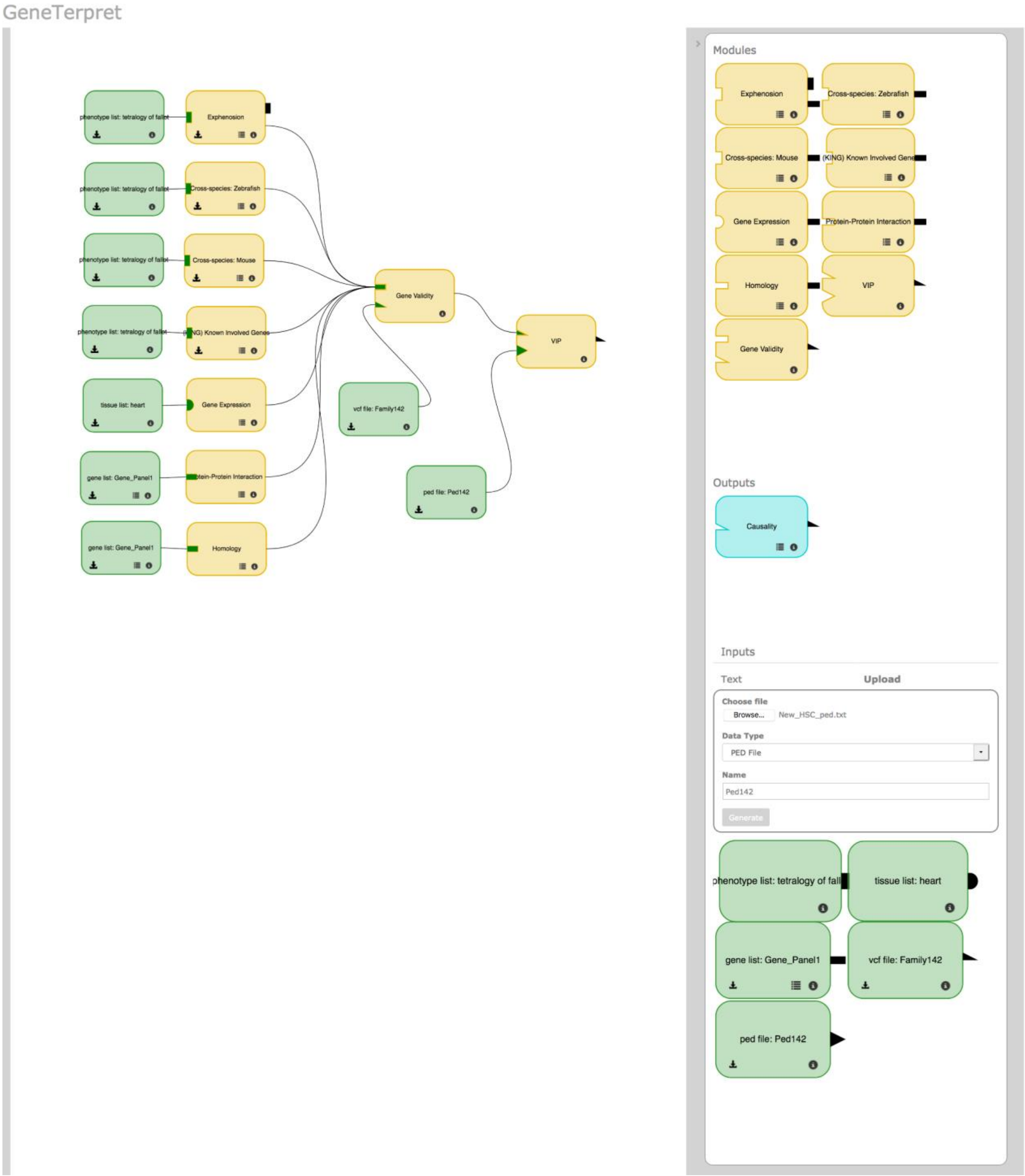


**Supplementary Figure S1) A snapshot of the *GeneTerpret* graphical user interface (*GeneTerpret GUI*).** A general interpretation routine is depicted as an example. The user selects the needed modules from the top right panel; then drags and drops them one by one in the left workspace panel. Furthermore, the tissue or phenotype/disease of interest can be directly entered by the user as an input in the bottom right panel and the generated module could be dragged and dropped in the left workspace panel. The users can upload their annotated VCF file, gene list(s), family information (PED file) and phenotypes/diseases list as further input for *GeneTerpret* by tapping on the upload tab in the bottom right panel and drag and drop the assigned generated module for the uploaded file in the workspace panel in the left side.

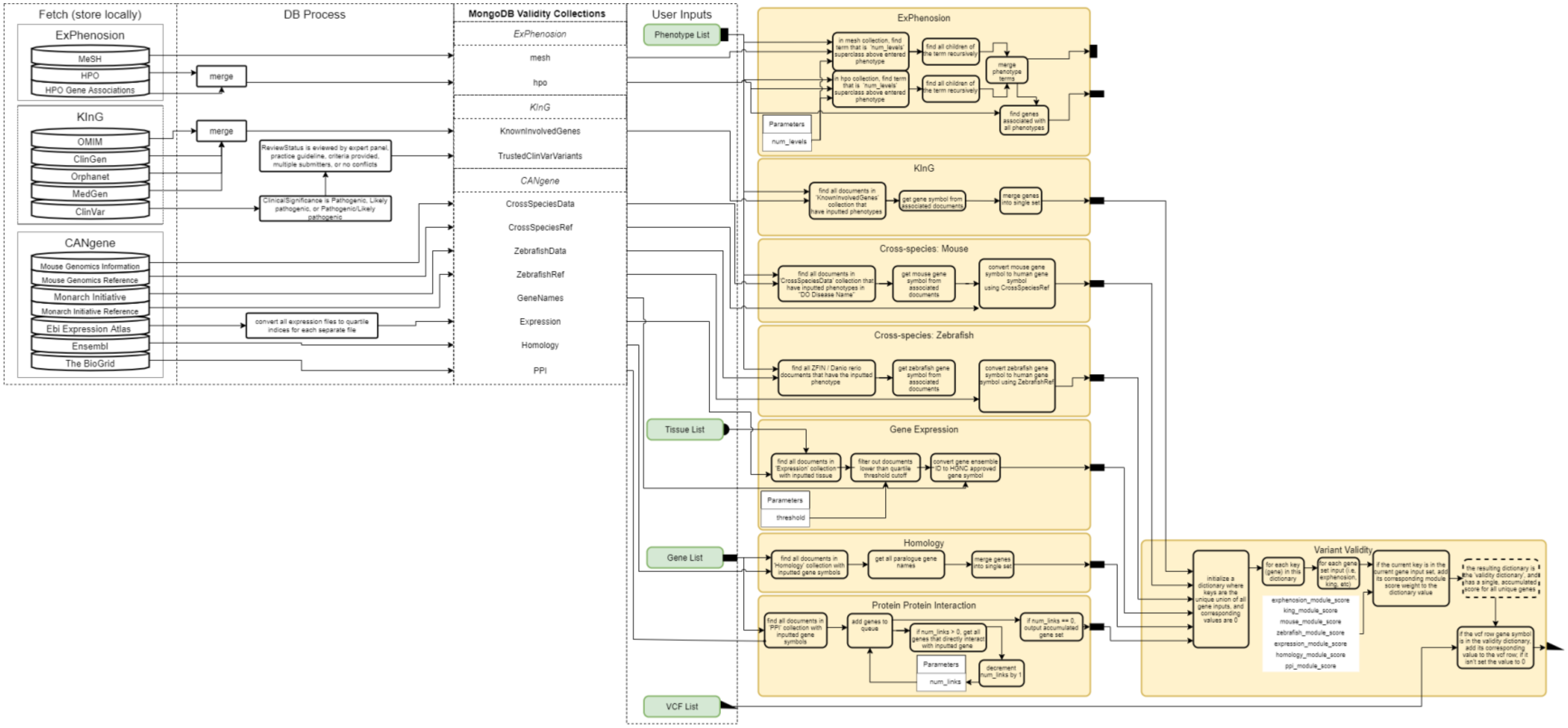


**Supplementary Figure S2) Gene Validity Module architecture.** External databases are first fetched and filtered based on certain criteria, and the results are entered into MongoDB collections. *ExPhenosion*, *CanGene*, and *KING* modules take in user input and the MongoDB collections to perform their functions.

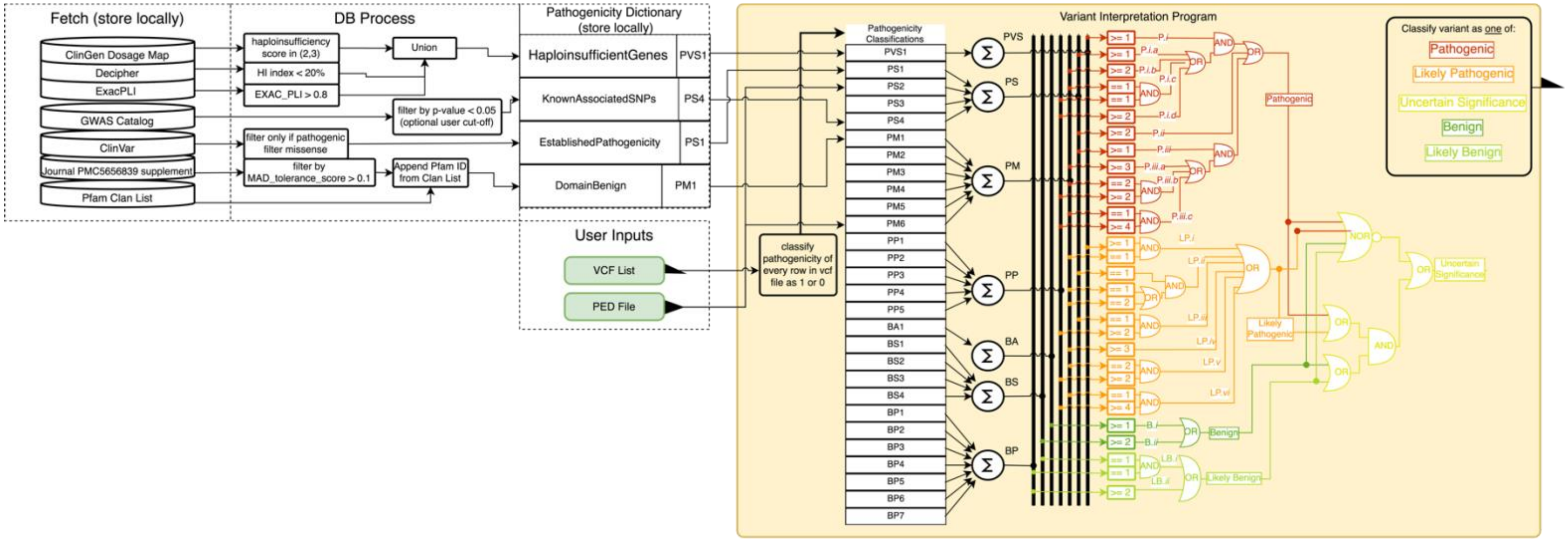


**Supplementary Figure S3) Variant Interpretation Program (*VIP*) internal structure.** External databases used for VIP are fetched and processed, with the output being stored in a MongoDB collection. In *VIP*, each database collection is associated with a specific *ACMG* classification, but not all classifications use these collections. Each row of the input VCF file is inputted to all classifications, flagging them as 1 or 0. Then, using the logic outlined in Table S2, the individual classifications are combined to provide the pathogenicity classification for each variant.

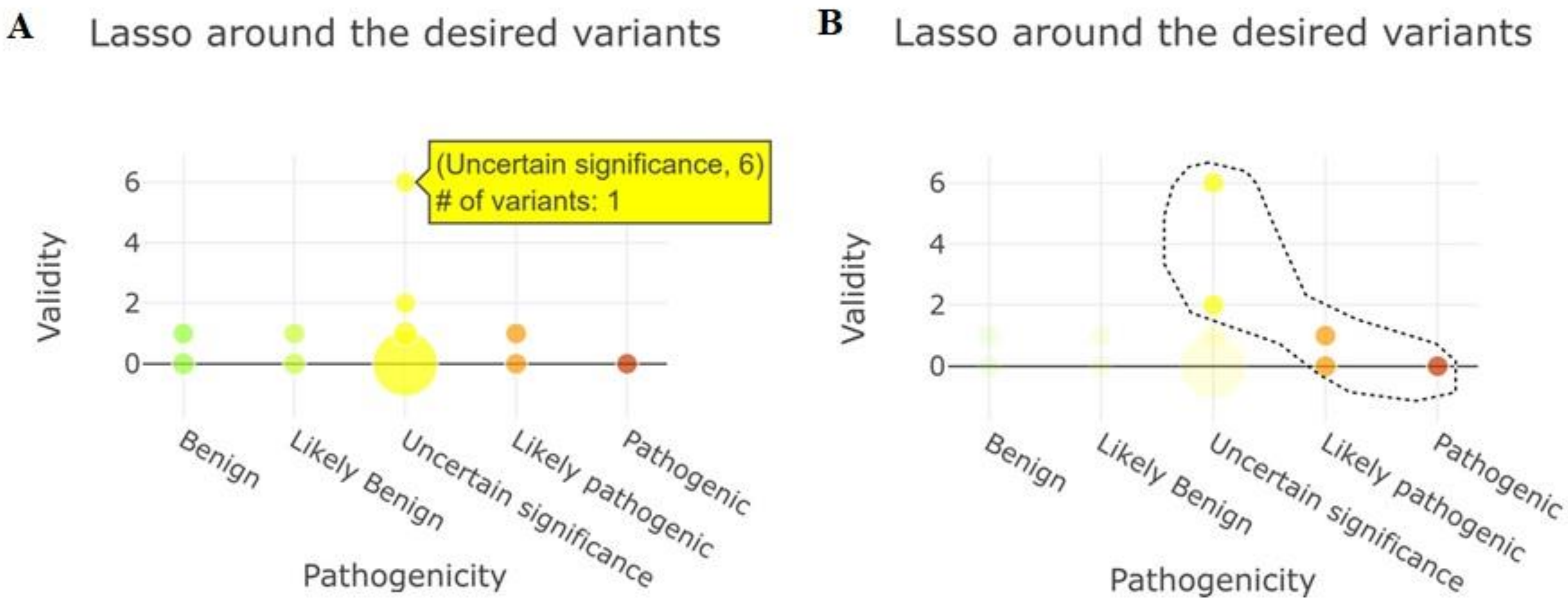


**Supplementary Figure S4) Overview of the causality module output**; **(A)** the interactive visualization of variant distribution in the validity-pathogenicity space allows users to explore the desired variants. Dark green, light green, yellow, orange, and red colours represent the pathogenicity of variants in a 5-tier system: benign, likely benign, uncertain significance, likely pathogenic, and pathogenic variants. **(B)** Lasso filter allows the analyst to select the desired variants and filter them to a downloadable VCF file.
